# Development and optimization of a *Barley stripe mosaic virus* (BSMV)-mediated gene editing system to improve Fusarium head blight (FHB) resistance in wheat

**DOI:** 10.1101/2022.02.25.481987

**Authors:** Hui Chen, Zhenqi Su, Bin Tian, Yang Liu, Yuhui Pang, Volodymyr Kavetskyi, Harold N. Trick, Guihua Bai

## Abstract

Fusarium head blight (FHB) is a devastating disease in wheat that causes million dollars of wheat yield losses annually in the U.S.A. Recently we demonstrated that wheat carry FHB susceptible genes and knocking out the susceptible gene increased FHB resistance, which suggests that manipulating susceptible genes using gene editing may open a new avenue to create new sources of FHB resistance. However, wheat genome editing uses gene transformation to deliver CRISPR/Cas9 and gRNA into plants, and most wheat genotypes are not suitable for transformation due to low rates of callus induction and regeneration, therefore it cannot be used in practical wheat breeding. In this study, we developed a new *Barley stripe mosaic virus* (BSMV)-mediated gRNA delivery system that does not need the gene transformation and tissue culture and it can be used in any genotypes for gene function validation and editing. We used this system edited the susceptible allele of *Fhb1*, a major FHB resistance gene. We demonstrated that the edited trait is heritable in different genetic backgrounds and knocking out Fhb1 susceptible allele improved FHB resistance in wheat. We also modified system to improve editing efficiency by using floral dip agroinfiltration and adding RNA mobility sequences to the gRNA in the viral vectors. We believe this work will facilitate wheat FHB resistance research and gene editing in cereal crops and will benefit cereal crop researchers and breeders worldwide.

---

Fusarium head blight (FHB) caused by *Fusarium graminearum* is a destructive wheat (*Triticum aestivum*) disease worldwide and growing FHB-resistant cultivars is an effective approach to minimize FHB damage in wheat (Bai *et al*., 2018). Among more than 600 quantitative trait loci (QTLs) reported for FHB resistance, only *Fhb1* showing a major effect on FHB resistance originated from wheat (Zheng *et al*., 2020). Therefore, it has been widely used in breeding programs worldwide (Bai *et al*., 2018). Previously, we cloned a histidine-rich calcium-binding protein gene (*TaHRC*) as the causal gene for *Fhb1* and demonstrated that a large deletion near the start codon of *TaHRC* reduced FHB susceptibility (Su *et al*., 2019), indicating that wheat carries susceptibility genes (S-genes) and knockout of these S-genes may enhance plant resistance. Recently, CRISPR/Cas9 genome editing system has been used to precisely knock out S-genes to generate resistant mutants in different crops (Chen *et al*., 2019); however, it requires transforming CRISPR/Cas9 and gRNA into plants using gene bombardment or *Agrobacterium*. Unfortunately, most wheat genotypes have extremely low callus induction and regeneration efficiency, which limits the application of genome editing in wheat breeding. Therefore, a new gRNA delivery system that bypasses the tissue culture is critical to the successful use of gene editing in routine wheat breeding.

*Barley stripe mosaic virus* (BSMV) has been an efficient vector used for virus-induced gene silencing and RNA interference in wheat (Yuan *et al*., 2011). Vectors have been engineered to deliver gRNAs into both dicots and monocots including wheat, and several genes have been edited in the leaves of these species (Hu *et al*., 2019). However, it remains unknown if the edited sequence changes are inheritable although BSMV can be transmitted to the next plant generation through germline cells (Hu *et al*., 2019). Here we developed a BSMV-mediated gRNA delivery system that bypasses the tissue culture for editing *TaHRC* in FHB-susceptible wheat cultivars (Chen *et al*., 2018).

We firstly overexpressed Cas9 (Cas9-OE) in Bobwhite using particle bombardment transformation, and then engineered a single guide RNA (sgRNA) into the downstream of the γb ORF in a BSMV RNA γ genome vector (Figure 1a). To evaluate its editing efficiency in wheat, we constructed sgRNAs for two wheat genes, phytoene desaturase (*TaPDS*) for albinism and *TaHRC* for FHB resistance, using this vector and then agroinfiltrated four-week-old *N. benthamiana* leaves with mixed *Agrobacterium* cultures harboring equal concentrations of the BSMVα, BSMVβ and BSMVγ::TaPDS, or BSMVα, BSMVβ and BSMVγ::TaHRC vectors. The inoculated tobacco leaves were checked for the presence of gRNA fragments using RT-PCR at six days post-inoculation (dpi) and then the sap was used to inoculate Cas9-OE Bobwhite wheat plants at the two-leaf stage (Figure 1b). At 21 dpi, the mutation efficiency (indel%) was evaluated for *TaPDS* (58%) and *TaHRC* (49%) in the systemically infected wheat leaves using T7 endonuclease I (T7E1) mutation detection assays and further confirmed by Sanger sequencing (Figure 1c, d). To evaluate the feasibility to use this system for multiplex gene editing, mixed *Agrobacterium* cultures harboring BSMVγ::TaPDS and BSMVγ::TaHRC vectors were co-agroinfiltrated into tobacco leaves, then the infected sap was used to inoculate the prepared Cas9-OE Bobwhite plants mentioned above. The multiplex gene editing system effectively delivered gRNA and precisely edited the target wheat gene although the mutation efficiency was slightly lower (41% for *TaPDS* and 47% for *TaHRC*) than those for singleplex gene editing (Figure 1c, d, e).

**Figure 1.**
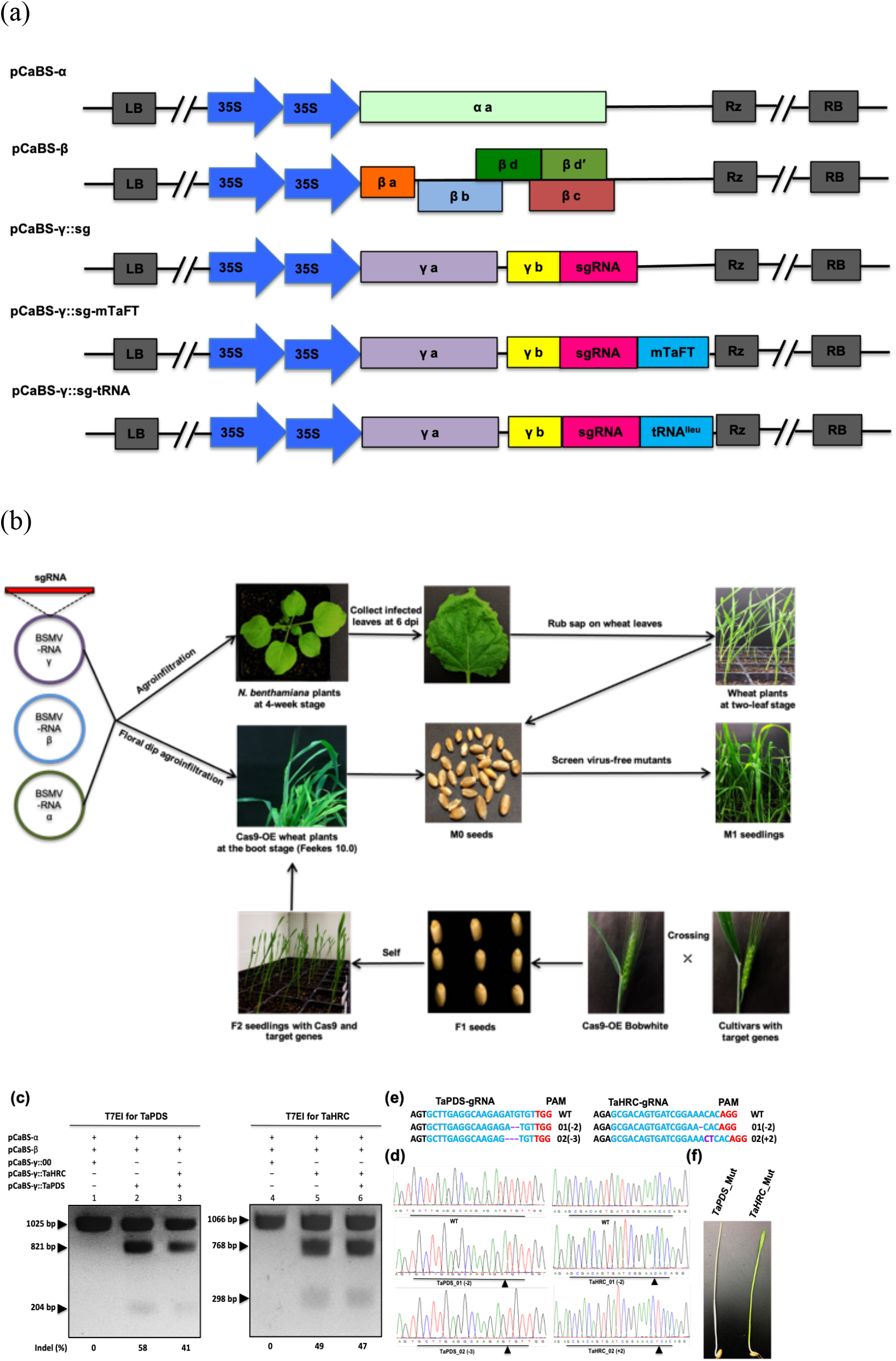
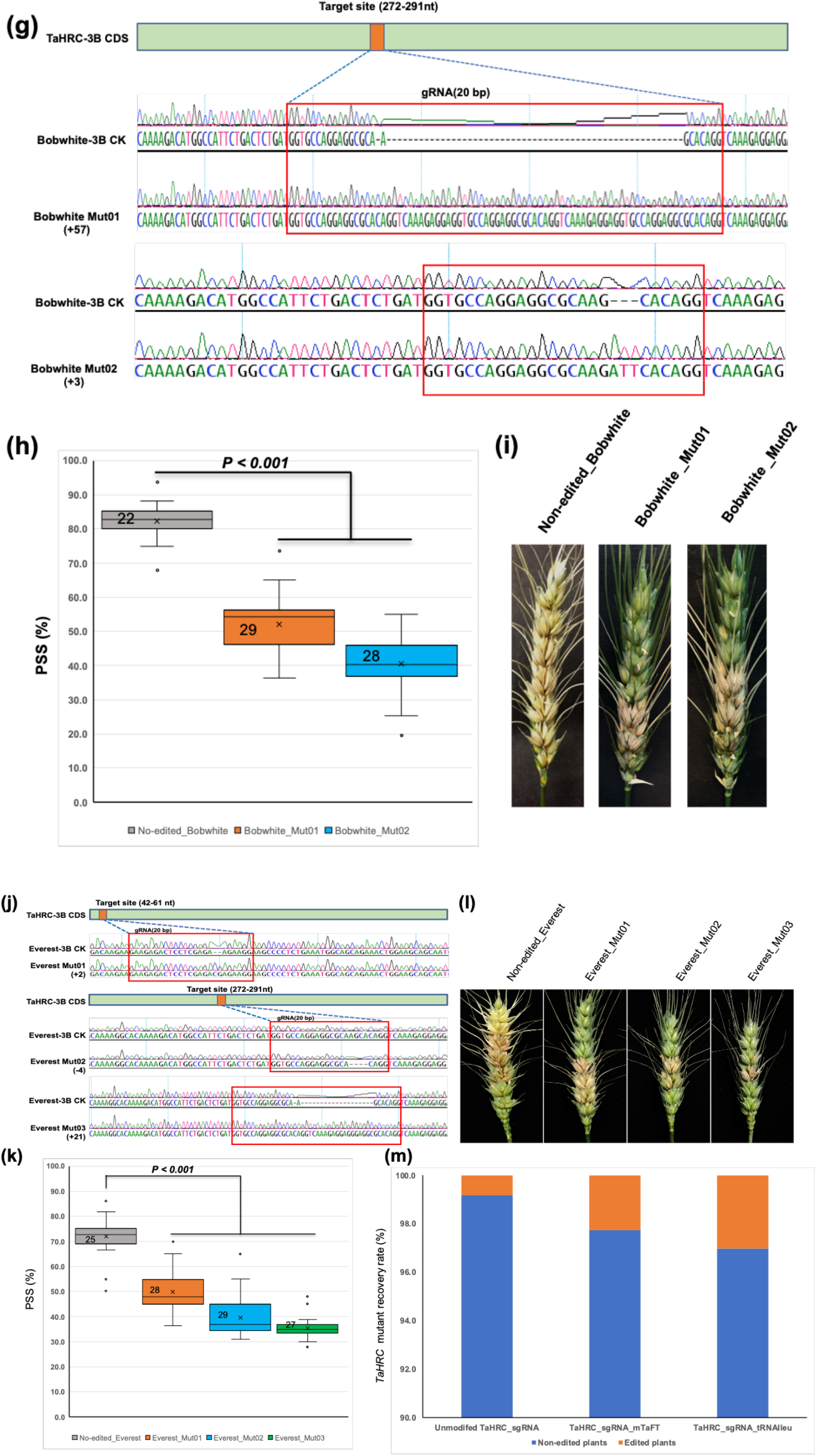
A BSMV-mediated gene editing system for improving wheat FHB resistance. (a) BSMV RNAα, RNAβ and RNAγ_sgRNA vectors used in wheat gene editing. (b) Workflow for the BSMV-mediated gene editing system. (c) T7EI assays detected *TaPDS* (1025 bp) and *TaHRC* (1066 bp) in BSMV infected Cas9-overexpressed wheat leaves. Lanes 1 and 4 are vector controls, lanes 2 and 3 show *TaPDS*-specific cleavage bands (821 and 204 bp), and lanes 5 and 6 show *TaHRC*-specific cleavage bands (768 and 298 bp) from the mutants. (d) and (e) Edited In/del sequences at *TaPDS* and *TaHRC* target sites of the mutants. (f) Seedlings of a *TaPDS* albino mutant (left) and *TaHRC* mutant. (g) Sequence of edited and non-edited *TaHRC* in Bobwhite showing insertions in Mut01 (57 bp) and Mut02 (3 bp). (h) and (i) FHB symptoms and percentages of symptomatic spikelets (PSS = 100 x infected spikelets /total spikelets in a spike) between the mutants and non-mutant control. (j) Sequences of edited and non-edited *TaHRC* in Everest. (k) and (l) PSS and infected spikes of mutants and non-edited control. (m) Recovery rates of *TaHRC* mutants generated using TaHRC_sgRNA, TaHRC_sgRNA-mTaFT and TaHRC_sgRNA-tRNA^Ileu^ vectors. In (de) and (e), 01 = Mut01 and 02 = Mut02. In (e) and (j), + and - refer to inserted and deleted nucleotides in bp, respectively. In (g) and (j), orange bars are target sites of gRNAs. In (h) and (k), boxes, whiskers, center line, crosses and numbers represent 25th-75th percentile, ranges, medians, means and sample size. *P*-values were from Student’s t-tests.

To determine if the edited sequences are heritable, we screened 187 M1 plants for *TaPDS* and 245 M1 plants for *TaHRC* and identified one *TaPDS* albino mutant and two *TaHRC* mutants (Figure 1f). To improve the edited mutant recovery rate, we used the floral-dip agroinfiltration method (Zale *et al*., 2009) to treat the spikes of Cas9-OE Bobwhite at preanthesis (Feekes 10.0) (Figure 1b), and obtained two mutants, Bobwhite_Mut01 and Bobwhite_Mut02 with 57- and 3-nucleotide insertions at *TaHRC*, respectively (Figure 1c, e, g). These mutants were phenotyped for FHB resistance in a growth chamber by injecting a conidial spore suspension of *F. graminearum* (GZ3639) into a central spikelet in a spike at early anthesis using a syringe (Su *et al*., 2019). The percentage of symptomatic spikelets (PSS) in a spike in the mutants was significantly lower than the non-edited control at 14 dpi (Figure 1h, i), demonstrating that the BSMV-mediated gene editing is heritable and knockout of the *TaHRC* susceptible allele improved wheat FHB resistance.

To expand the utility of this editing system in other wheat cultivars with low transformation efficiency, we transferred the *Cas9* gene into a locally adapted winter wheat cultivar ‘Everest’ by crossing Everest to a Cas9-OE Bobwhite plant and selecting Cas9-OE Everest F_2_ progeny that carried homozygous *Cas9* and the target gene. The immature spikes of the selected Cas9-OE Everest at preanthesis (Feekes 10.0) were agroinfiltrated with the mixed *Agrobacterium* cultures harboring the BSMVα, BSMVβ and BSMVγ::TaHRC vectors using the floral dip method. Sequencing 318 M1 plants at the target site identified three Everest-*TaHRC* mutants with a 2bp insertion (+2) in Everest_Mut01, a 4bp deletion (−4) in Everest_Mut02 and a 21bp insertion (+21) in Everest_Mut03 (Figure 1j). FHB phenotyping in a growth chamber showed a significantly lower PSS in the mutants than in the non-edited Everest (Figure 1k, l), confirming that loss-of-function mutations in *TaHRC* increased FHB resistance in the Everest derivatives. Therefore, the BSMV-mediated gRNA delivery system has potential to create new sources of FHB resistance and can be used for gene function validation in gene cloning experiments. The newly developed Everest derivative mutants with the *Fhb1* resistance allele are BSMV and transgene free and can potentially be further used as a resistant parent in wheat breeding.

Several recent studies demonstrated that edited mutant recovery rate can be increased by adding endogenous mobile RNA sequences such as *Flowering Locus T* (*FT*) and transferring RNA-like sequences (tRNAs) to the 3-end of sgRNAs. These modifications can improve accessibility into plant shoot apical meristems (SAM) and therefore increase frequencies of heritable sequence changes in the edited genes and mutant recovery rates in the progeny (Ellison *et al*., 2020). To test this in the BSMV-mediated gene editing system, we fused a truncated wheat *FT* RNA sequence (mTaFT) with 135 bp of wheat Vrn3-a allele and a tRNA^Ileu^ mobile sequence (isoleucine) to the 3’-end of the sgRNA in the BSMV RNA γ genome vector that targeted the *TaHRC* in the Cas9-OE wheat plants (Figure 1a). We obtained a higher editing rate of *TaHRC* mutant lines from the M1 progeny (2.3% and 3.0%) using the modified TaHRC_sgRNA_mTaFT and TaHRC_sgRNA_tRNA^Ileu^ constructs than those (0.8%) using the TaHRC_sgRNA construct alone (Figure 1m). These results indicate that adding RNA mobility sequences in the virus vector increased the heritable sequence mutations in the edited genes and mutant recovery rate in wheat, concurring with Ellison et al. (2020), but disagreeing with Li et al. (2021). Different lengths or types of RNA mobile elements fused to the sgRNA in BSMV vector might contribute to the discrepancy.

In summary, we developed a BSMV-mediated gRNA delivery system coupled with floral dip agroinfiltration and gRNA sequence modification, which improved editing efficiency and successfully edited *TaHRC* in Bobwhite and Everest. We demonstrated that the new gene editing system has potential for application in routine breeding to improve wheat resistance FHB and other diseases without genotype limitation.

## Acknowledgement

This is contribution number 22-051-J from the Kansas Agricultural Experiment Station. We thank Dr. Dawei Li for providing the BSMV clones. This project was partially supported by the US Wheat and Barley Scab Initiative and the USDA National Research Initiative Competitive Grants 2017-67007-25939.

## Author contributions

HC and GB designed the project and wrote the manuscript, HC, ZS, BT, YL, YP, HT and VK performed experiments. All authors read and approved the manuscript.

